# Expansion microscopy with trypsin digestion and tyramide signal amplification (TT-ExM) for protein and lipid staining

**DOI:** 10.1101/2023.03.20.533392

**Authors:** Ueh-Ting Tim Wang, Xuejiao Tian, Yae-Huei Liou, Sue-Ping Lee, Chieh-Han Lu, Peilin Chen, Bi-Chang Cheb

**Affiliations:** Affiliated Senior High School of National Taiwan Normal University, Taipei, 10658; Research Center for Applied Sciences, Academia Sinica, Taipei 11529, Taiwan; Nano Science and Technology Program, Taiwan International Graduate Program, Academia Sinica 115, Taiwan; Department of Engineering and System Science, National Tsing Hua University, Hsinchu 300, Taiwan; Institute of Molecular Biology, Academia Sinica, Taipei, 11529, Taiwan, ROC

**Author notes:** To whom correspondence should be addressed. (B.C.C.).

**Keywords:** expansion microscopy, trypsin, tyramide signal amplification, lipid stain, mitochondrial DNA

## Abstract

Expansion microscopy, whereby the relative positions of biomolecules are physically increased via hydrogel expansion, can be used to reveal ultrafine structures of cells under a conventional microscope. Despite its utility for achieving super-resolution imaging, expansion microscopy suffers two major drawbacks, namely proteolysis and swelling effects that, respectively, induce protein loss and dilute fluorescence signals. Here, we report two improvements to expansion microscopy that overcome these two challenges, i.e., deploying trypsin digestion to reduce protein loss and tyramide signal amplification to enhance fluorescence signal. We name our new methodology TT-ExM to indicate dual trypsin and tyramide treatments. TT-ExM may be applied for both antibody and lipid staining. Notably, we demonstrate better protein retention for endoplasmic reticulum and mitochondrial markers in COS-7 cell cultures following 2-h trypsin treatment. Subsequent lipid staining revealed the complex 3D membrane structures in entire cells. Through combined lipid and DNA staining, our TT-ExM methodology highlighted mitochondria by revealing their DNA and membrane structures in cytoplasm, as well as the lipid-rich structures formed via phase separation in nuclei at interphase. We also observed lipid-rich chromosome matrices in the mitotic cells. Thus, TT-ExM significantly enhances fluorescent signals and generates high-quality and ultrafine-resolution images under confocal microscopy.

## 1. Introduction

Fluorescence microscopy is a powerful approach for visualizing the fine cellular structures of organisms because of its high sensitivity and specificity. By means of antibody staining or fluorescent protein tagging, proteins in cells can be localized according to the distributions of emitted signals from fluorophores. However, the spatial resolution of fluorescence microscopy is limited to approximately half the wavelength of fluorescence signals, termed the Abbe limit, thereby preventing visualizations of ultrafine subcellular structures. With the invention of super-resolution techniques—such as stimulated emission depletion microscopy (STED), structured illumination microscopy (SIM), photoactivated localization microscopy (PALM), and stochastic optical reconstruction microscopy (STORM)—it is now possible to image biological samples and circumvent the Abbe limit. Instead of improving optics to attain greater spatial resolution in super-resolution microscopy, expansion microscopy (ExM) was invented to bypass the optical diffraction limit by physically expanding biological samples in a hydrogel, enabling the effective resolution to be reduced to the nanometer scale under a conventional optical microscope [1-4].

Originally, expansion microscopy was designed so that samples were expanded 4-fold in one dimension [1], which was called x4 expansion microscopy. Later, x10 expansion microscopy was developed that expanded the samples 10-fold in one dimension [5, 6]. The effective resolution was therefore improved from 200 nm to 50 nm (for x4 expansion microscopy) or to 20 nm (for x10 expansion microscopy). This simple approach to super-resolution microscopy opened up the possibility of observing molecules and/or cellular structures at the nanoscale level without the need for sophisticated microscopes. Since various antibodies and staining reagents can be readily applied in expansion microscopy techniques, it can be used to monitor cell and tissue ultrastructures. For example, conventional antibodies and fluorescent protein labeling can be used to label cellular proteins[1-3, 6], and metabolic labeling [7] or membrane-binding fluorophore-cysteine-lysine-palmitoyl group (mCLING) technology [8, 9] have been developed for visualizing membranes. Notably, mCLING labels plasma membranes, so it can also be taken up during endocytosis for intracellular vesicle labeling [8]. In addition, DNA dyes can be used to label chromatin [10].

Despite expansion microscopy having already proven valuable in studies of the ultrafine structures of cells and tissues, it has some caveats. One obvious problem is the reduction in fluorescence signals inherent to diluting fluorophore concentrations. As the volume of a sample increases to at least 64- or 1000-fold (for x4 and x10 expansion, respectively), fluorophore concentrations concomitantly are diluted to 1/64 or 1/1000, which significantly reduces detectable fluorescence signals and impairs image quality, especially for 3D rendering. This dilution effect on fluorophore intensity is one of the major drawbacks of expansion microscopy. Another challenge with this technique is associated with sample preparation. To allow biological samples to expand isotropically with the hydrogel, the protein molecules in the samples have to be covalently anchored to the hydrogel before being subjected to protease-mediated digestion [1-6]. The anchored fluorophores can then freely move with the hydrogel upon its expansion in water. Proteinase K is routinely used to digest such samples since it is a potent protease and thus guarantees proteolytic efficiency [1-6]. However, proteinase K exhibits broad substrate specificity, recognizing both aliphatic and aromatic amino acids, so it digests protein samples into very small peptides. These latter may not be anchored to hydrogels and so can be lost during expansion. Thus, proteolysis also reduces fluorescence signals after sample expansion.

To enhance fluorophore signals and ensure their retention in expanded hydrogels, biotin-streptavidin complex [11] and trifunctional anchoring reagent [12] treatments have been incorporated into expansion microscopy methodologies to label protein molecules. Here, we have established a modified protocol using both tyramide signal amplification to increase overall fluorescence signals and trypsin digestion to enhance fluorophore retention. We call this new protocol expansion microscopy with trypsin digestion and tyramide signal amplification, i.e., TT-ExM in short. TT-ExM can be applied to both protein and lipid labeling, together with DNA counter-staining. The protocol is simple and easy to apply, yet it significantly increases fluorescence signals and retains fluorophores in the hydrogel-expanded samples. Entire 3D cell images can be readily recorded using a confocal laser scanner, resulting in at least a 4-fold increase in resolution.

## 2. Results and Discussion

### 2.1 The basic principle of TT-ExM

In order to enhance fluorescent signals in expansion microscopy, we made two modifications to the original protocol, as highlighted in red in the outline of TT-ExM depicted in Figure 1A. First, we used the Tyramide Signal Amplification (TSA) system to overcome the dilution effect caused by hydrogel-mediated expansion, with horseradish peroxidase (HRP)-conjugated secondary antibody replacing the typical fluorophore-conjugated secondary antibody. In the presence of peroxidase and hydrogen peroxide, Alexa fluor-555-conjugated tyramide can be covalently linked to the side chain of tyrosine [13], which enhances fluorescence signals (Figure 1B). For biotinylated molecules, the avidin/biotin-HRP complex (ABC) can be applied to recognize and covalently conjugate tyramide to molecules or nearby proteins. For membrane and lipid staining, we used biotin-DHPE to label the membrane and lipid-containing structures in cells. Since ABC effectively recruits HRP molecules to biotinylated molecules, the signal enhancement attainable through ABC is much greater than for HRP-conjugated secondary antibodies (Figure 1C). Thus, in general, the HRP-conjugated secondary antibody can be replaced with a combination of biotinylated secondary antibody and ABC [14].

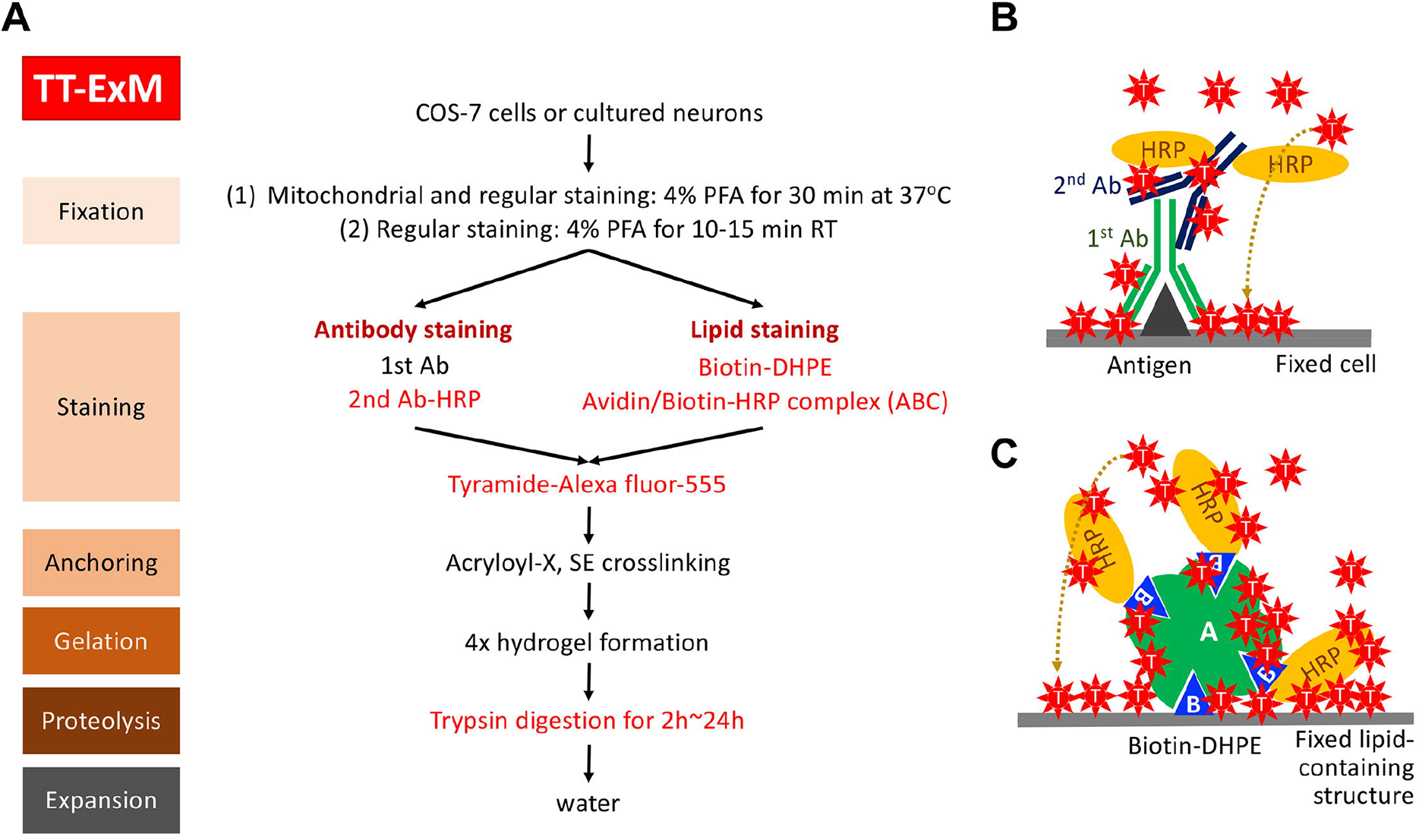

The second modification we did was to replace proteinase K with trypsin for proteolysis after the gelation step (Figure 1A). Proteinase K is a broad-specificity protease, which cuts at the carboxyl-terminal end of all aliphatic or aromatic amino acids (Table 1). Since the average frequency of all aliphatic or aromatic amino acids in protein molecules is 46.6% (Table 1), the average length of the proteolytic products using proteinase K is ∼2.1 amino acid residues, which greatly reduces the possibility of anchoring fluorophore-conjugated peptides to the hydrogel and thus retain them there during expansion. In contrast to proteinase K, trypsin cuts specifically at the carboxyl-terminal end of lysine and arginine. The combined usage frequency of lysine and arginine in proteins is ∼11.4% (Table 1). Therefore, the average length of the proteolytic products following trypsinization is ∼8.8 amino acids. Accordingly, we anticipated that trypsinization would prove more efficient in terms of fluorophore retention in expansion microscopy.

**Table 1.** Comparison of proteinase K and trypsin digestion.

|  | proteinase K | trypsin |
| --- | --- | --- |
| Concentration of the protease | 8 units/ml | 0.015% |
| Ingredients of the digestion buffer | Tris, pH 8.0, 50 mM;<br>EDTA, 1 mM;<br>Triton-X-100, 0.5%;<br>*Guanidine HCl, 0.8 M | Tris, pH 8.0, 35 mM;<br>EDTA, 0.7 mM;<br>Triton-X-100, 0.35% |
| Cleavage sites of the proteases | Aliphatic or aromatic amino acids, including leucine (7.6%), alanine (7.4%), valine (6.8%), isoleucine (3.8%), glycine (7.4%), phenylalanine (4%), proline (5%), tryptophan (1.3%) or tyrosine | Lysine (7.2%) or arginine (4.2%)** |
|  | (3.3%)** |  |

\* Guanidine HCl is a chaotropic agent, which can destroy intramolecular and intermolecular hydrogen bonds, thereby unraveling the three-dimensional structure of proteins and nucleic acids, which increases the hydrolytic efficiency of proteases. This chemical was only used in the proteinase K-based hydrolysis reaction, and not in the trypsin reaction.
\*\* The percentages shown in parentheses after the amino acids represent the average frequency of use of each amino acid in vertebrates [21], thereby reflecting the number of times that the protease cleaves a protein molecule with a length of 100 amino acids under ideal conditions.

### 2.2 A combination of TSA and trypsinization retains convincing fluorescent signals in ExM

To evaluate the improvements achieved using TSA and trypsinization, first we tested antibodies that recognize protein disulfide isomerase (PDI, a marker of endoplasmic reticulum (ER)), and cytochrome c oxidase subunit 5B (COX5B, a mitochondrial marker), in TT-ExM. To prepare the hydrogel, we followed the protocol for 4x ExM [1], but with some modifications [15]. The size of the gel was expanded ∼4.8-fold (i.e., (14 wells + 15 wells)/2/3 wells=4.8; Figure 2A). Next, we examined immunoreactivities to confirm the successful application of the TSA system before hydrogel expansion (Figure 2B). After trypsinization and expansion, a piece of hydrogel was removed and poststained with the DNA dye PicoGreen. For this experiment, a 20X objective lens was used to acquire fluorescent signals before and after expansion. From the resulting images, trypsinized COS-7 cells and their nuclei clearly extend evenly throughout the hydrogel (Figure 2C, 2D). Note that the fields of view in Figures 2B, 2C and 2D are the same, in which labeled cells were expanded ∼4-fold. Importantly, PDI immunoreactivity signals were well preserved and the images were easily acquired using an inverted LSM700 confocal laser-scanning microscope (Figure 2C). Although COX5B signals were weaker than those of PDI upon expansion, we still acquired reasonably strong signals (Figure 2D).

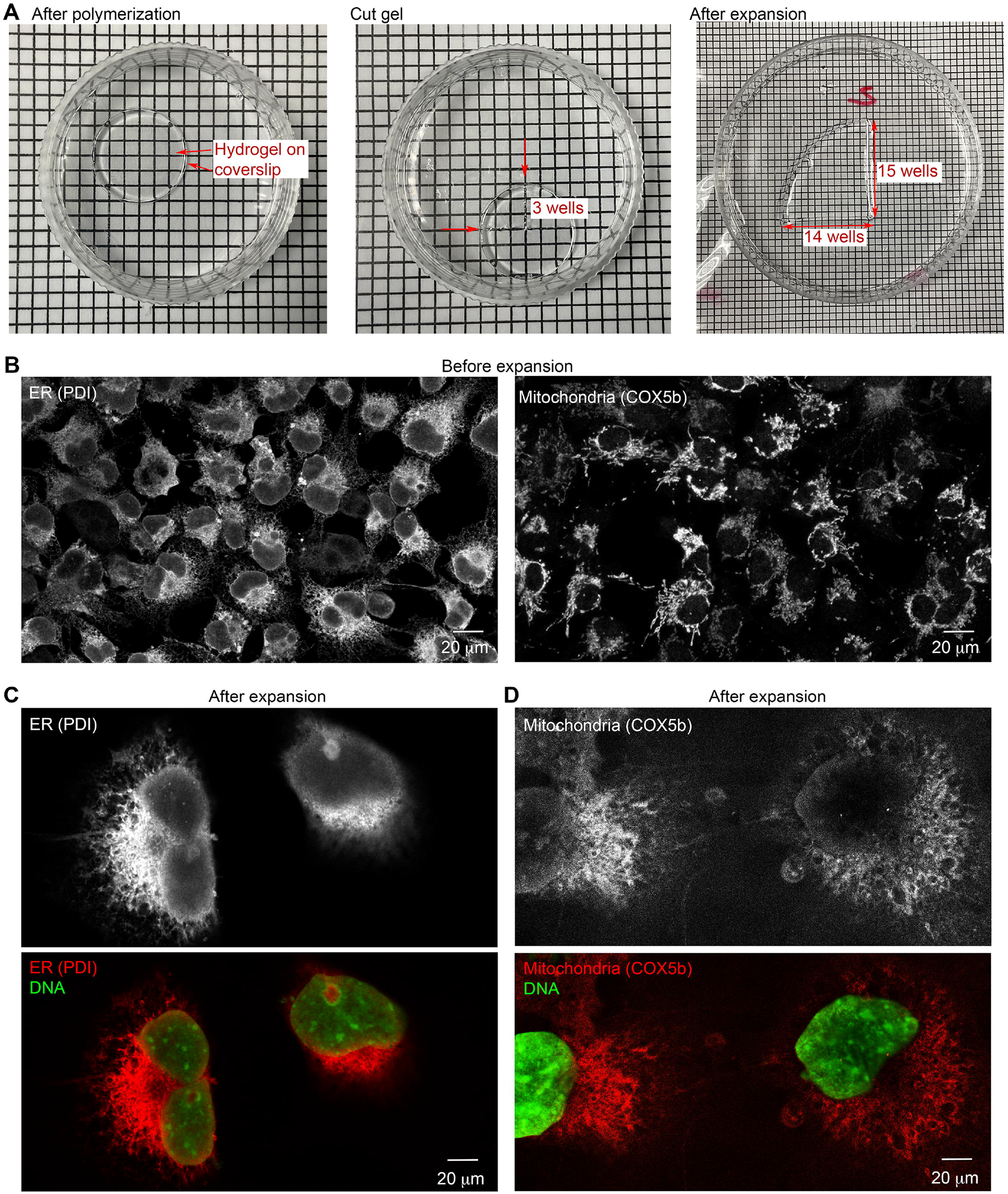

To evaluate our protocol further, we conducted a comparison of trypsin and proteinase K proteolysis treatments. We found that the immunoreactivity of proteinase K-treated cells was much weaker than for trypsinized COS-7 cells under the same experimental conditions (Figure 3A). To compare the images, we needed to increase the PDI fluorescence intensity of proteinase K-treated cells 3-fold relative to trypsin-treated cells, indicating that fluorescence signals were indeed more strongly retained. We observed the mesh-like structure of ER (Figure 3A, right) and some bagel-shaped mitochondria in our images achieved using only a 20X objective lens (Figure 3B). When we enhanced the signals of PicoGreen-labeled nuclei to near saturation, we observed tiny puncta of PicoGreen signal in the cytoplasm. These signals primarily overlapped with or were adjacent to COX5B immunoreactivities (Figure 3C), thus potentially representing mitochondrial DNA. Based on these results, we have demonstrated that our TT-ExM protocol noticeably enhances and retains fluorescence signals in hydrogels and that clear images can be easily acquired using a 20X objective lens under an inverted confocal microscope.

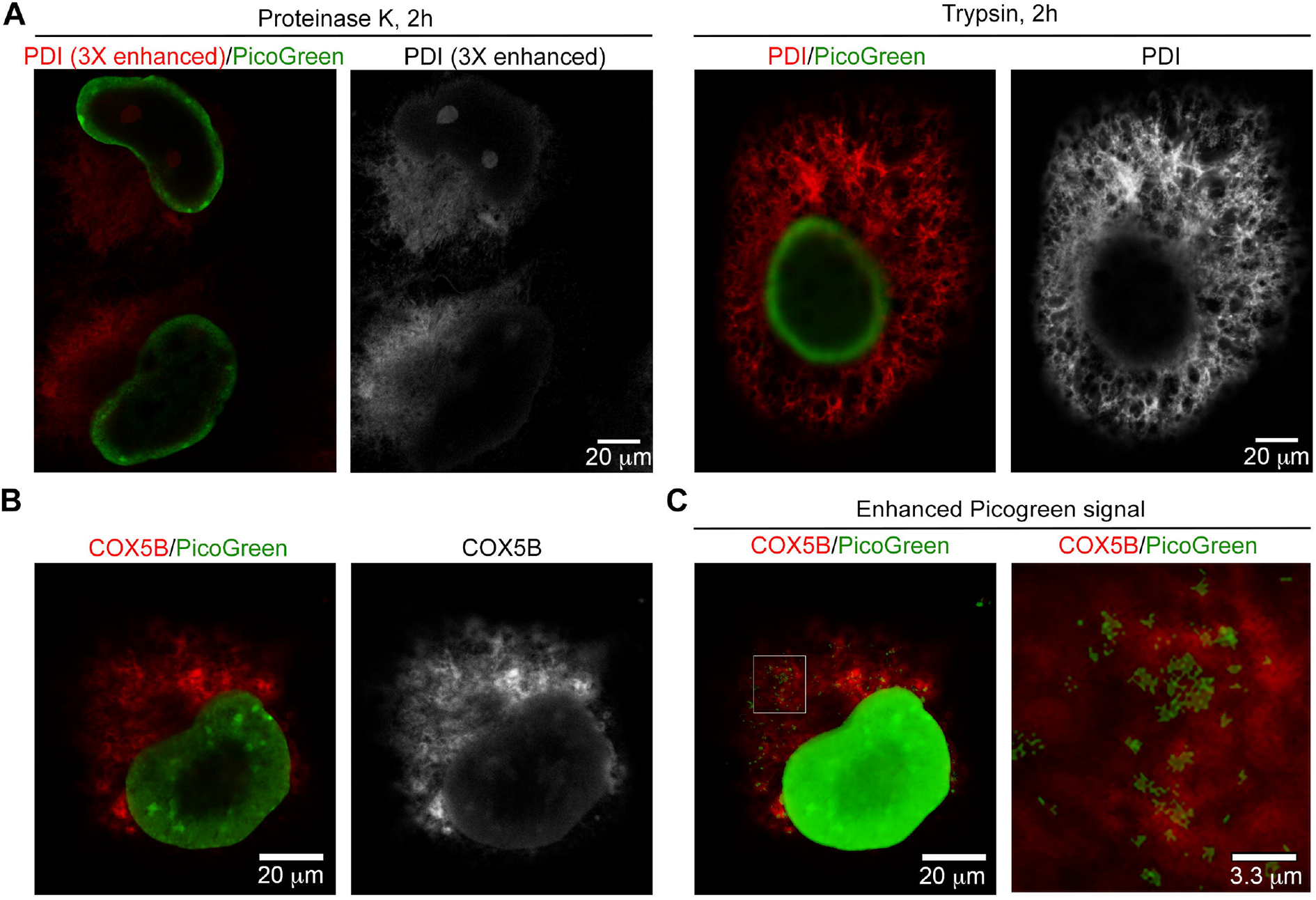

### 2.3 Application of TT-ExM for lipid staining

Apart from applications in imaging protein distributions, it can also be important to visualize cellular structures by means of lipid staining. Biotin-DHPE can be used to label lipid-containing structures [16-18], so we investigated if it can be applied to TT-ExM to stain the lipid structures of cells. We dissolved biotin-DHPE in chloroform and then diluted the mix with 70% or 50% ethanol for cell staining, with incubation times of 150 min or overnight. Among these four conditions, we found that the strongest biotin-DHPE fluorescent signals were obtained using 50% ethanol dilution and overnight incubation (Figure 4A). Thus, we adopted this combination for subsequent experiments. Using a 20X objective lens, not only could we clearly visualize plasma membrane, but biotin-DHPE also labeled the intracellular membrane structures of COS-7 cells (Figure 4A). To acquire images with a higher spatial resolution, we deployed an LSM980 confocal laser-scanning microscope with AiryScan2 and observed the nucleus and cytoplasmic mesh-like structures (presumably ER) in a cell at interphase using biotin-DHPE staining (Figure 4B, left and middle). Intriguingly, we noticed that biotin-DHPE labeled the entire nucleus, including the nuclear matrix and even some nuclear aggregates lacking DNA (Figure 4B, left and middle). We also performed sectioning in the z-plane and reconstructed a 3D image of an entire cell (except for the nucleus) using Imaris, which revealed the complex membrane organization of the entire cell (Figure 4B, right). Our images depicted certain COS-7 cells undergoing cell division, with representative examples in metaphase and anaphase shown in Figures 4C and 4D, respectively. Significantly, the biotin-DHPE signal was enriched between the condensed chromosomes of the dividing cells (Figure 4C, 4D), with these special biotin-DHPE-stained structures being attached to chromosomes (Figure 4C, 4D, right panel), which is very similar to the pattern generated by staining for the cell proliferation marker Ki-67 [19, 20].

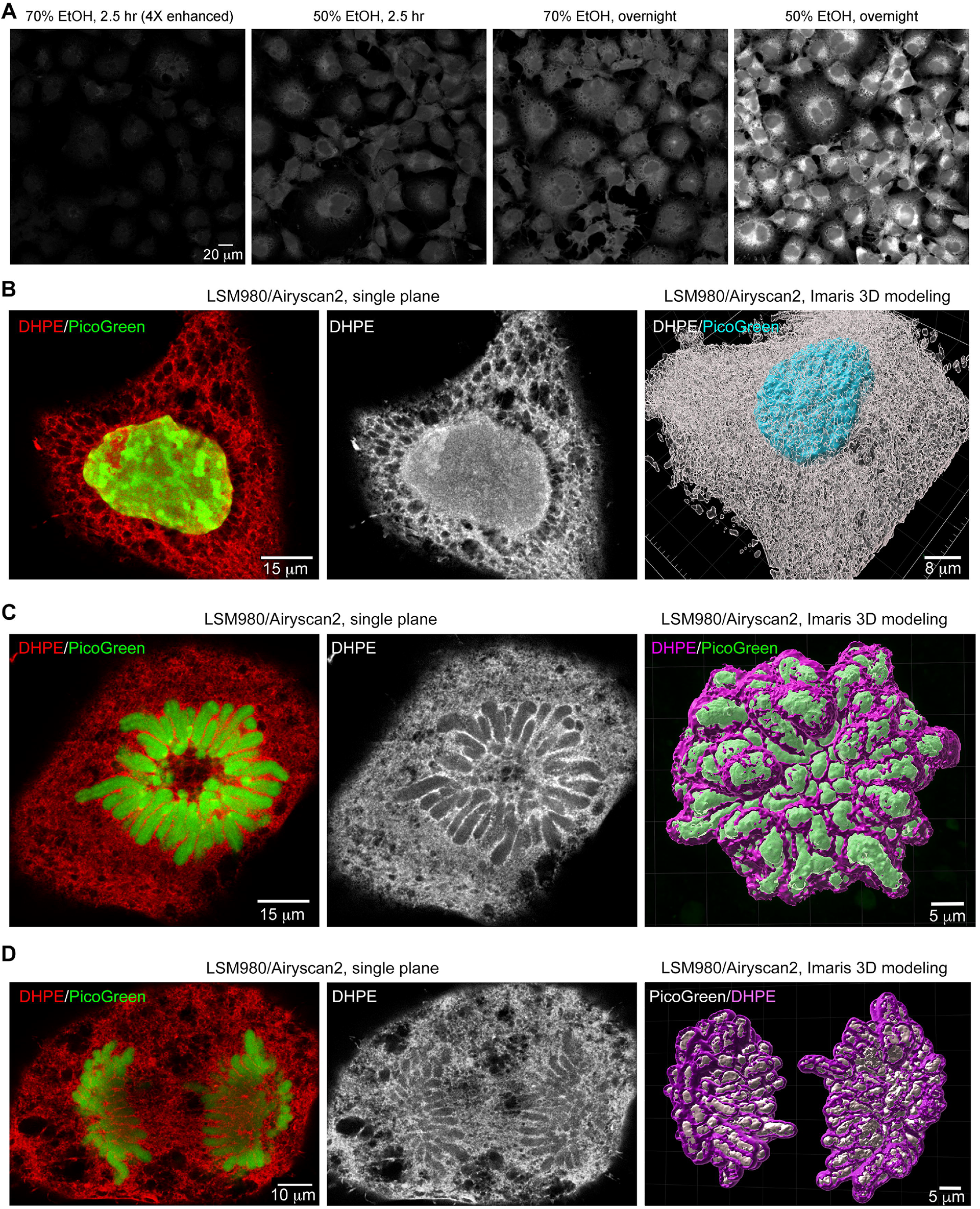

Therefore, we term this biotin-DHPE-stained structure as the “chromosome matrix”, which may be involved in chromosome organization and integrity during cell division.

Based on these images, we show that a combination of biotin-DHPE and TSA can be used in TT-ExM to label entire COS-7 cells, revealing many fine cellular details. Moreover, we have been able to observe the ultrafine structures of the cells under confocal microscopy.

### 2.4 Observing mitochondria and nuclei in cultured neurons by means of TT-ExM

In addition to studying COS-7 cells, we also endeavored to image primary cultured neurons prepared from mouse brains using TT-ExM and biotin-DHPE staining (Figure 5). Neuronal cell bodies are approximately 3-fold smaller than COS-7 cells, so we only deployed LSM980 confocal microscopy with AiryScan2 to visualize neurons. We found that biotin-DHPE clearly stained the entire neurons, including their soma, dendrites and axons (Figure 5A, before expansion). After hydrogel-based expansion, we clearly observed numerous DHPE-dense tubular structures in the cultured neurons (Figure 5B, after expansion). Since these tiny tubular structures were ∼0.5-1 µm in width and frequently harbored small PicoGreen-positive puncta (Figure 5C), we speculate they represent mitochondria in the cultured neurons. Thus, TT-ExM deployed with lipid and DNA stainings can enable visualization of mitochondria in neurons.

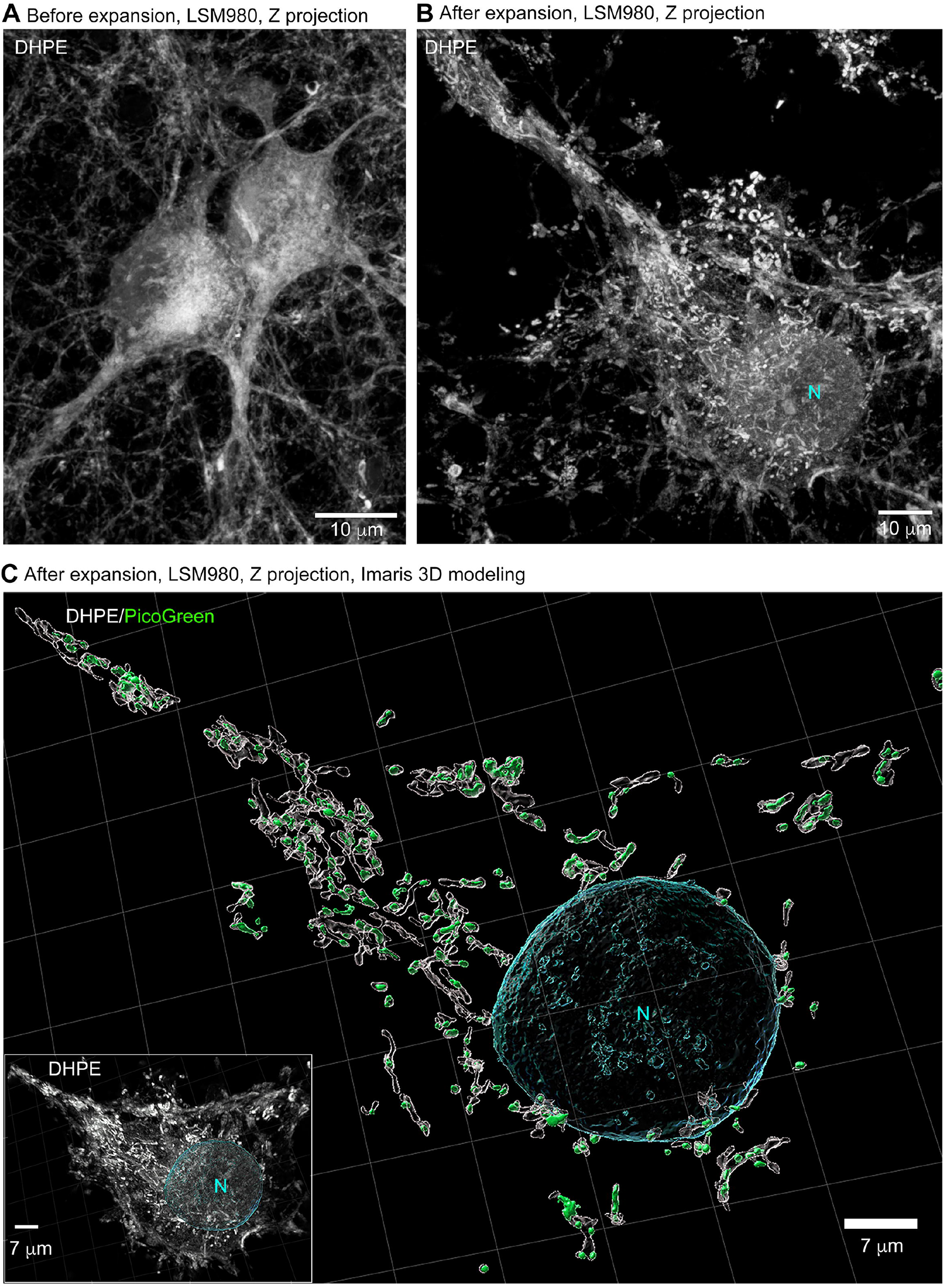

As for COS-7 cells, biotin-DHPE staining also highlighted the nuclear structure within neurons (Figure 6A). Moreover, biotin-DHPE staining revealed numerous tiny droplets and some large condensates in the nucleus (Figure 6A, 6B, pink arrows), which tended to be separate from but adjacent to chromosomal DNA (Figure 6A, 6B). In particular, the large DHPE-positive condensates were always adjacent to large DNA aggregates, presumably representing the nucleolar structure (Figure 6B, white arrows). In postmitotic neurons, DHPE-stained nuclear structures seemed to contribute to organizing chromosomal DNA and gene expression, unexpectedly revealing a special organization of lipid-containing structures in the nucleus.

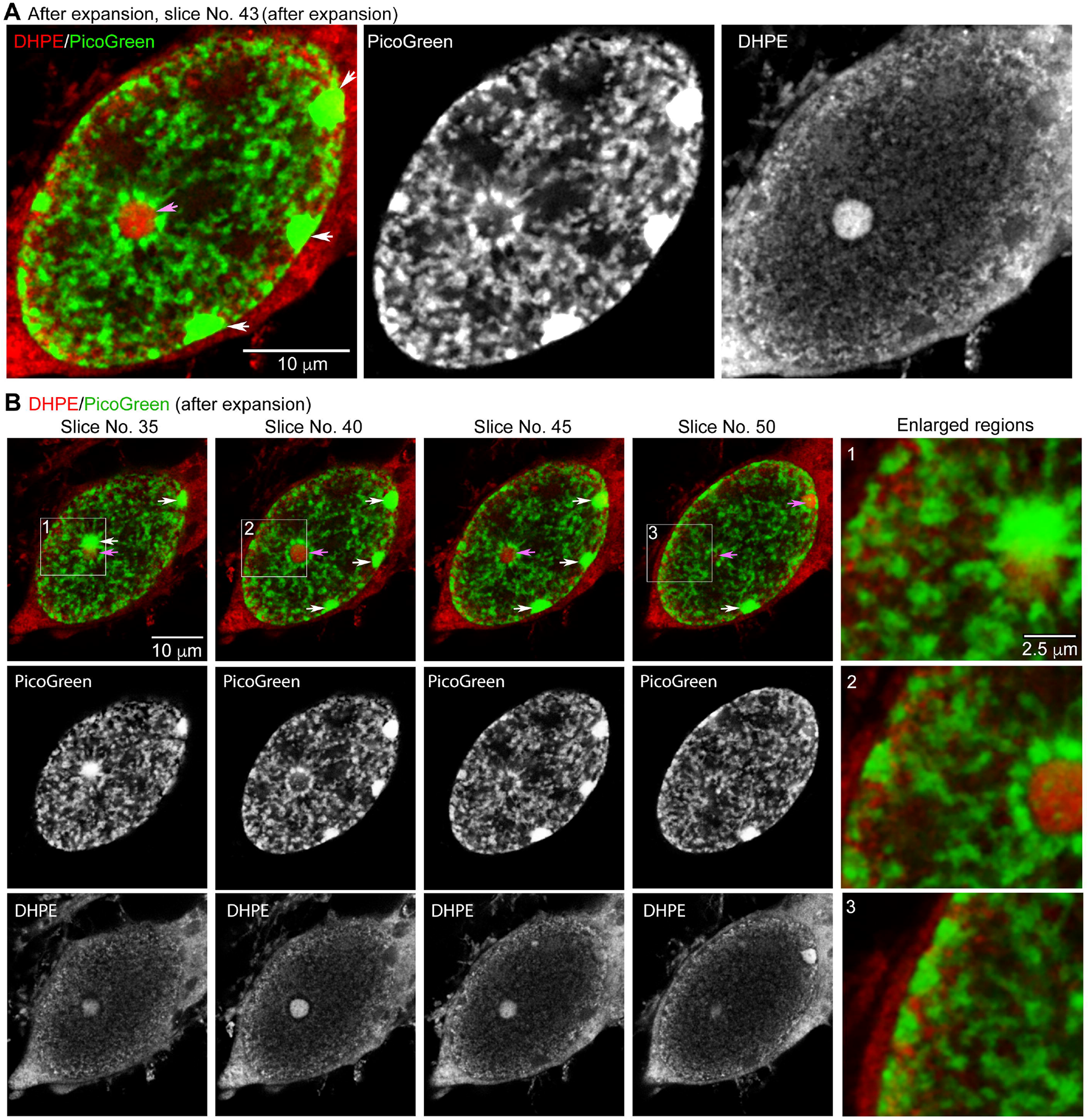

## 3. Conclusions

In summary, herein we present an improved protocol for expansion microscopy (TT-EXM) that enhances signal retention of protein and lipid cellular components and amplifies fluorescence signals. We adopted trypsin digestion for proteolysis of cultured cells, which engenders fewer cutting sites than the proteinase K treatment used in the original ExM method, resulting in more peptide fragments of organelles such as ER or mitochondria being anchored to the hydrogel matrix of TT-ExM. In addition, we deployed TSA coupled with HRP-conjugated secondary antibody to recruit more fluorophores, increasing fluorescence signal intensity. Moreover, by applying a combination of biotin-DHPE membrane dye, avidin/biotin-HRP complex and tyramide amplification, we were easily able to observe lipid-containing cellular components under a regular microscope, circumventing on oft-cited issue for previous expansion microscopy protocols due to their detergent-treatment steps. Through biotin-DHPE labeling, we uncovered in 3D a chromosome matrix at the chromosomal boundary during cell division. Therefore, TT-ExM also represents a convenient approach for investigating the nanoscale-level distributions of lipid-rich compartments in 3D during phase separation of biological systems by means of light microscopy.

Overall then, we have successfully applied TT-ExM to cultured cell systems for super-resolution imaging. By combining TT-ExM with DHPE staining, we have also achieved expansion microscopy for lipid visualization, which could potentially be useful for hydrogel-based expansion of large tissues, such as Drosophila melanogaster brain or zebrafish organs.

## 4. Material and Methods

### 4.1 Reagents and material for cell culture

Dulbecco’s Modified Eagle Medium (DMEM), Gibco, 11965-084; Fetal bovine serum (FBS), HyClone, SH30084.03; L-Glutamine, Gibco, A29168-01; Penicillin/Streptomycin, Gibco, 15140-122; 10X PBS, pH 7.4, Lonza, 17-517Q, 10-fold dilution before use; 0.05% Trypsin-EDTA, Gibco, 25300-054; poly-L-lysine, Sigma-Aldrich, P26360, 1 mg/ml; No. 1 coverslip, 15-mm in diameter, Fisher, 1942-10015.

### 4.2 Cell preparation

The COS-7 cell line (ATCC, CRL-1651), an African green monkey kidney fibroblast-like cell line, was maintained in DMEM supplemented with 10% FBS, 1% penicillin/streptomycin and 2 mM glutamine in a 5% CO2 incubator at 37 °C. To prepare cell samples for expansion microscopy, COS-7 cells that grew to 100% confluency in a 10-cm culture dish were harvested and resuspended in 10 ml DMEM. The cell suspension (0.4 ml) was added into one well of a 12-well plate containing DMEM growth medium (1 ml/well) with a poly-L-lysine-coated coverslip. The cells were then placed back in a 5% CO2 incubator at 37 °C. One day later, cells were ready for immunostaining. Hippocampal and cortical mixed cultures were kindly provided by Dr. Yi-Ping Hsueh’s laboratory.

### 4.3 Reagents for immunofluorescence staining

Paraformaldehyde (PFA), Merck, 1.04005.1000; Triton X-100, Sigma-Aldrich, T8787; Tris, VRW, 0826; Blocking reagent for immunostaining, Tyramide Signal Amplification kit, PerkinElmer, 581428; Cytochrome c oxidase subunit 5B (COX5B) rabbit polyclonal antibody, Proteintech, 11418-2-AP, 3-6 µg/ml; Protein disulfide isomerase (PDI) rabbit polyclonal antibody, Proteintech, 11245-1-AP, 1-2 µg/ml; Beta-tubulin mouse monoclonal antibody, Sigma-Aldrich, T4026, 6 µg/ml; Horseradish peroxidase (HRP)-conjugated anti-rabbit IgG, secondary antibody, Jackson ImmunoResearch Laboratory, 711-035-152, 1:500 dilution; Alexa fluor-488-conjugated anti-mouse IgG, secondary antibody, ThermoFisher, A28175, 1:500 dilution; Alexa fluor-555-conjugated anti-rabbit IgG, secondary antibody, ThermoFisher, A32732, 1:500 dilution; Alexa fluor-555-conjugated tyramide, Invitrogen, B40955, 1:100 dilution; Amplification diluent, PerkinElmer, FP1050; PicoGreen, ThermoFisher, P11495, 1:1000 dilution.

### 4.4 Immunostaining with tyramide signal amplification

COS-7 cells that grew on coverslips were first washed with PBS three times and fixed with 4% PFA in PBS for 15 min for regular staining. Alternatively, cells were washed three times with PBS pre-warmed to 37 °C and then fixed with 4% PFA in PBS pre-warmed to 37 °C for 30 min at 37 °C to observe mitochondria in cells. After washing twice with Tris-buffered saline (TBS), TBST (TBS + 0.3% Tritin-X-100) was added, before incubating for 10 min for permeabilization. After rinsing once with TBST, 0.5% Blocking solution was added for 30 min. Cells were then incubated with the primary antibodies diluted in blocking solution for 2 h at room temperature or overnight at 4 °C. After washing three times with TBST, secondary antibodies were added and incubated for 1.5 h at room temperature. After rinsing three times with TBST to remove unbound antibodies, Alexa fluor-555-conjugated tyramide was diluted 100-fold with amplification diluent and applied to cells for 10-20 min. The cells were then rinsed three times with TBST to stop the reaction. Before crosslinking, immunofluorescence signals were checked under an inverted fluorescence microscope to confirm successful staining.

### 4.5 Reagents for lipid staining

Paraformaldehyde (PFA), Merck, 1.04005.1000; 1x PBS; N-(Biotinoyl)-1,2-dihexadecanoyl-sn-glycero-3-phosphoethanolamine, triethylammonium salt (Biotin-DHPE), Biotium, 60022; VECTASTAIN ABC-HRP Kit, peroxidase (standard), Vector Laboratory, PK-4000; Alexa fluor-555-conjugated tyramide, Invitrogen, B40955, 1:100 dilution; Amplification diluent, PerkinElmer, FP1050; PicoGreen, ThermoFisher, P11495, 1:1000 dilution.

### 4.6 Lipid staining using tyramide signal amplification

Biotin-DHPE was dissolved in chloroform at a concentration of 10 mg/ml. After washing three times with PBS, COS-7 cells or cultured neurons were fixed with 4% PFA in PBS for 15 min at room temperature. To observe mitochondria, the PBS and 4% PFA were pre-warmed to 37 °C directly before use. After washing three times with TBS, fixed cells were transferred to 70% or 50% ethanol containing 0.1 mg/ml biotin-DHPE and incubated for 2.5 h or overnight. An equal amount of reagents A and B of an ABC-HRP Kit was mixed in TBS at a dilution of 1:50 for at least 30 min. The premixed ABC solution was then added to biotin-DHPE-labeled cells for 1 h. After washing, Alexa fluor-555-conjugated tyramide was diluted 1:100 in amplification diluent and applied to cells for 15 min. After extensive washing, the stained cells were then ready for anchoring to a hydrogel.

### 4.7 Reagents for anchoring and making the acrylamide hydrogel

6-((acryloyl)amino)hexanoic acid (Acryloyl-X, SE), ThermoFisher, A-20770; Dimethyl sulfoxide (DMSO), Thermo Fisher, D12345; Sodium acrylate (SA), Sigma-Aldrich, 408220; Acrylamide (AA), Sigma-Aldrich, A9099; N, N’-methylenebisacrylamide (MBAA), ThermoFisher, 1551624;

Ammonium persulfate (APS), Sigma-Aldrich, A9164; N,N,N,,N,-Tetramethylethylenediamine (TEMED), Sigma-Aldrich, T7024; 4-Hydroxy-TEMPO, ALDRICH, 176141; Sodium Chloride, J.T.Baker, 3624-69.

### 4.8 Anchoring

Before gelation, the stained cells were incubated with acryloyl-X, SE to crosslink protein molecules with the acrylamide gel. Acryloyl-X, SE was first dissolved in DMSO at a concentration of 10 mg/ml and then diluted to 0.1 mg/ml in PBS right before adding it to the stained cells. The samples were incubated overnight at 4 °C for reaction. Directly before gelation, the samples were washed three times with PBS to remove excess Acryloyl-X, SE.

### 4.9 Gelation

To make sufficient acrylamide solution for six gel discs of 0.5 mm thickness and 15 mm diameter, 564 µl acrylamide monomer solution (containing 8.6% sodium acrylate, 2.5% acrylamide, 0.15% N,N’-methylenebisacrylamide and 11.7% sodium chloride in PBS) was mixed with 12 µl 10% TEMED, 6 µl 1% TEMPO, 6 µl water and 12 µl 10% APS. After extensively vortexing, 88 µl of the mixture was immediately added onto parafilm, which was placed in a 3.5-cm dish. After removing the extra PBS, a coverslip was placed on top of the acrylamide mixture with the side hosting cells facing down. Gelation was achieved at 37 °C for 2 h.

### 4.10 Enzyme digestion and expansion of the hydrogel

Approximately a quarter of the hydrogel disc was cut with a scalpel and measured using graph paper. The gel was then gently picked up with a flat brush and placed in a solution containing a protease for hydrolysis. We tested two proteases, i.e., proteinase K and trypsin, incubated either for 2 h or overnight. The concentrations of these two enzymes, the composition of the digestion buffer, and their cleavage sites are listed in Table 1. The digested hydrogel was completely immersed in double-distilled water in a 10-cm petridish. After at least 30 min, the double-distilled water was replaced with fresh water and this process was repeated at least three times. The size of the swollen hydrogel was again measured using graph paper.

### 4.11 Poststaining with PicoGreen DNA dye

The expanded hydrogel was placed in water containing the DNA dye PicoGreen (1:1000) for at least 2 h. To completely penetrate into nuclei, PicoGreen staining longer than overnight was required.

### 4.12 Imaging using a LSM700 confocal laser scanner and image processing

An appropriate size (∼0.5 × 1 cm) of the expanded hydrogel containing COS-7 cells was placed on a 24×50 mm coverslip with the side hosting cells facing down. The excess water was carefully removed. The expanded samples were then observed under an inverted confocal microscope LSM700 (Zeiss) with an Plan Apo 20X/NA=0.8 DICII objective lens. Non-expanded samples were directly mounted and observed using the same 20X objective lens under the same microscope for comparison with expanded samples. Fluorescence images were captured with Zen software 2009 (https://www.zeiss.com/microscopy/en/products/software.html) from the Zeiss Corporation. Single focal plane images were collected and then analyzed using ImageJ/Fiji software (V 2.1.0/1.53c, https://imagej.nih.gov/ij/index.html). The images were then converted into a format recognizable by general image software and assembled using Adobe Photoshop 2020.

Imaging using a confocal laser scanner LSM980 system with Airyscan2 and image processing using Labkit and Imaris

For cultured neurons, the expanded gel was placed in a water-filled mold on a 0.17-mm-thick coverslip. Images were recorded using a LSM980 laser scanning confocal microscope equipped with multiplex mode SR-4Y (2X sampling) and an Airyscan 2 detector. A LD LCI Plan-Apochromate 40X/1.2 Imm Korr DIC M27 objective was adopted for image acquisition. We applied 488 nm and 561 nm laser excitations for DNA and lipid signals, respectively. The 1844×1844 pixel images in the Z series were acquired at 0.2 µm intervals. The Z-series images were further processed to conduct 3D surface construction using Imaris 9.9.1 software with the machine learning Fiji-Labkit plugin for segmentation. The shortest distance and smooth options were chosen as Imaris parameter settings.

The channel of interest was then directed to Labkit. In Labkit, the pencil tool was used to draw on images and identify “foreground” and “background” regions. This segmentation was sent back to Imaris to create the surface. In neurons, mitochondrion-like surfaces were defined by tubular lipid signals with a minimal object-object distance to DNA signals. The images were then converted into a format recognizable by general image software and assembled using Adobe Photoshop 2020.

## Author Contributions

U.T.T.W. and X.T. planned and performed the expansion experiments. Y.H.L and S.P.L. performed imaging experiments and image processing. U.T.T.W. performed immunostaining experiments. C.H.L and P.C. were involved in manuscript preparation. B.C.C. planned and managed the project.

## Acknowledgments

We thank Dr. Hsiao-Tang Hu at the Institute of Molecular Biology, Academia Sinica, for providing hippocampal and cortical mixed cultures. The authors would like to acknowledge financial support from the National Science and Technology Council of Taiwan (MOST 109-2628-M-001-001-MY4 (B.C.C.) and Academia Sinica of Taiwan [(AS-CDA-107-M08 (B.C.C.), AS-iMATE-110-43 (B.C.C.)].

